# The spatial organization of ascending auditory pathway microstructural maturation from infancy through adolescence using a novel fiber tracking approach

**DOI:** 10.1101/2024.06.10.597798

**Authors:** Kirsten M. Lynch, Stefanie C. Bodison, Ryan P. Cabeen, Arthur W. Toga, Courtney C.J. Voelker

**Affiliations:** Laboratory of Neuro Imaging (LONI), USC Mark and Mary Stevens Institute for Neuroimaging and Informatics, USC Keck School of Medicine, Los Angeles, CA, USA; Department of Occupational Therapy, College of Public Health and Health Professions, University of Florida, Gainesville, FL, USA; Otology/Neurotology, Pacific Neuroscience Institute, Los Angeles, CA

**Keywords:** Auditory pathway, child development, microstructure, tractography, NODDI, white matter

## Abstract

Auditory perception is established through experience-dependent stimuli exposure during sensitive developmental periods; however, little is known regarding the structural development of the central auditory pathway in humans. The present study characterized the regional developmental trajectories of the ascending auditory pathway from the brainstem to the auditory cortex from infancy through adolescence using a novel diffusion MRI-based tractography approach and along-tract analyses. We used diffusion tensor imaging (DTI) and neurite orientation dispersion and density imaging (NODDI) to quantify the magnitude and timing of auditory pathway microstructural maturation. We found spatially varying patterns of white matter maturation along the length of the tract, with inferior brainstem regions developing earlier than thalamocortical projections and left hemisphere tracts developing earlier than the right. These results help to characterize the processes that give rise to functional auditory processing and may provide a baseline for detecting abnormal development.

**Highlights:**

- We characterize the microstructural maturation of the auditory pathway
- We show structural development of the auditory pathway is dynamic and heterogeneous
- Diffusion metrics AD, RD and MD reach adult-like levels earlier than FA and NDI
- Brainstem microstructure matures earlier than the subcortical white matter
- Maturation of the right auditory pathway continues later than the left

## 1. Introduction

The auditory system is a fundamental sensory perceptual system that enables the identification and localization of sounds in the external environment. Auditory perception is established through exposure to experience-dependent stimuli during critical developmental periods of heightened plasticity in early life (Gabard-Durnam and McLaughlin, 2020) and plays a critical role in cognitive development (Ortiz-Mantilla et al., 2019). Infants born with congenital sensorineural hearing loss face many obstacles in the acquisition of speech later in life (Kral et al., 2019); however, cochlear implantation early in life during sensitive developmental periods can restore auditory function and enable normal acquisition of spoken language (Nicholas and Geers, 2013; Niparko et al., 2010; Robbins et al., 2004). Given the important role of myelinating processes on the closure of critical periods of auditory plasticity (Kalish et al., 2020; Mcgee et al., 2005), characterization of auditory white matter maturation is an important step towards understanding typical and atypical child auditory development, and possibly guiding interventional treatment planning. However, few studies have investigated the neural microstructure of the pathways that facilitate transmission of auditory information to the brain during typical development using neuroimaging techniques.

The ascending auditory pathway conveys spectrotemporal auditory information from the receptors in the inner ear to the primary auditory cortex (Peterson et al., 2021). The neural pathways that support this information transfer are complex and involve polysynaptic relays of decussating fibers through the brainstem, midbrain, thalamus and subcortical white matter (**Figure 1A**). In summary, the vestibulocochlear nerve (cranial nerve VIII) carries auditory information from the organ of Corti within the cochlea to the cochlear nucleus (CN) in the brainstem through the internal auditory canal. Fibers from the CN then bifurcate and synapse in the ispilateral and contralateral superior olivary nuclei (SON) in the trapezoid body of the brainstem where binaural cues essential for auditory localization are processed (Yin and Chan, 1990). Auditory information is then conveyed from the bilateral SON via the lateral lemniscus to the inferior colliculus (IC) (De Martino et al., 2013). Fibers then project to the moderating body of the medial geniculate nucleus (MGN) of the thalamus, and then pass through the acoustic radiation and terminate in the primary auditory cortex (A1) located within the superior temporal gyrus (Westerhausen et al., 2009). The auditory pathway undergoes a comparatively protracted developmental process that differs markedly from the early maturation observed in other primary sensory systems, such as the visual and somatosensory systems (Kinney et al., 1988; Pujol et al., 2006; Yakovlev and Lecours, 1967), presumably to accommodate language acquisition and cognitive development during childhood. This extended developmental time course therefore renders the auditory system particularly susceptible to auditory deficits during sensitive windows of enhanced plasticity, as auditory deprivation in infants and young children disrupts a critical window for receptive language development (Gabard-Durnam and McLaughlin, 2020).

**Figure 1.**
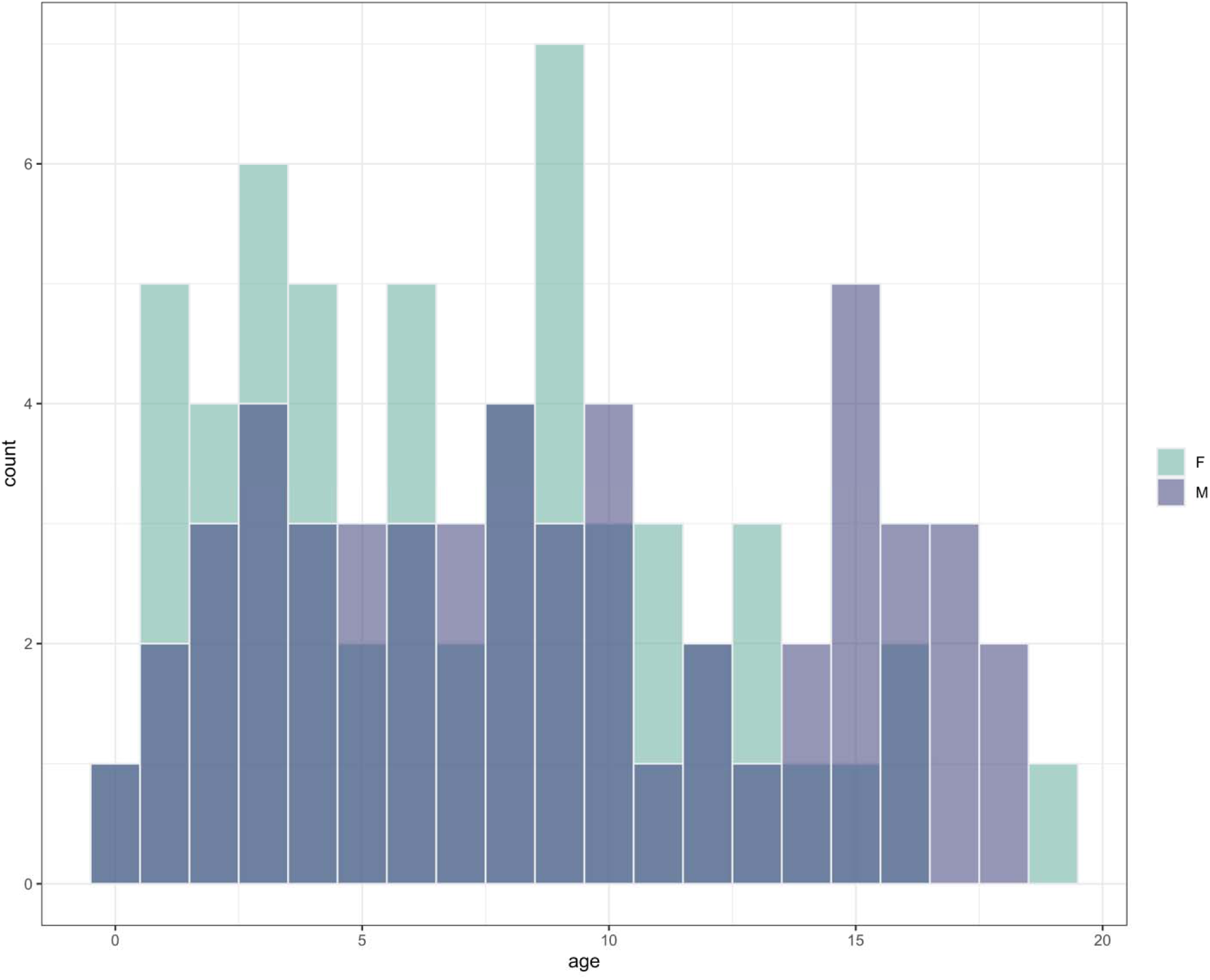
Age distribution of subjects included in the study stratified by sex. F = female; M = male.

Diffusion MRI (dMRI) is a non-invasive neuroimaging technique that is sensitive to the displacement patterns of diffusing water molecules and provides unique insight into characteristics of *in vivo* white matter structure (Beaulieu, 2009). Quantitative models derived from dMRI contrasts, such as diffusion tensor imaging (DTI) and neurite orientation dispersion and density imaging (NODDI), provide unique insight into the integrity of white matter microstructure. DTI is a dMRI model sensitive to white matter cellular features that influence diffusion anisotropy and tissue water content. The DTI metrics fractional anisotropy (FA), mean diffusivity (MD) and radial diffusivity (RD) may reflect axon caliber, myelination and axonal density (Barazany et al., 2009; Beaulieu, 2002; Song et al., 2002), while axial diffusivity (AD) may be associated with axonal injury and degradation (Concha et al., 2006; Song et al., 2003). While the metrics derived from the DTI model are sensitive to developmental processes, they are not specific and the relative contribution of different cellular features to changes in DTI metrics is unclear. NODDI is a multi-compartment diffusion model that estimates tissue compartments based on the restrictive nature of water diffusion patterns using multi-shell diffusion data (Zhang et al., 2012). The neurite density index (NDI), which models the fraction of a voxel occupied by axons, provides a more specific measure of axonal density and accounts for more age-related variance during child development compared to the DTI metric fractional anisotropy (FA) (Genc et al., 2017). Orientation dispersion index (ODI) provides an index of orientational complexity indicative of fiber dispersion. Therefore, utilization of metrics derived from both DTI and NODDI can provide complementary insight into different developmental processes within the auditory pathway.

The developmental time course of major white matter tract microstructure using DTI and NODDI metrics have been well studied in pediatric populations, which broadly show dynamic and spatially-varying patterns of maturation with age (Lebel et al., 2019a; Lebel and Deoni, 2018). However, few studies have assessed the microstructural development of the auditory pathway in vivo. In fact, the majority of research on the development of the auditory pathways and their associated cortical functions have been conducted in children with (Berman et al., 2016)varying degrees of hearing loss (Huang et al., 2015; Wang et al., 2019; Wu et al., 2016b) and are restricted to tractography of the auditory pathway thalamocortical projections (Berman et al., 2016; Wu et al., 2016a). Research into the structural connectivity of the auditory pathway *in vivo*, particularly within the brainstem, is complicated by a number of factors. Brainstem anatomy is characterized by a mixture of white matter bundles and subcortical nuclei, and dMRI tractography performance is inhibited by the polysynaptic nature of the auditory pathway (Mori and Tournier, 2013; Wakana et al., 2004). In order to mitigate the inherent difficulties of delineating the auditory pathway from the brainstem to the auditory cortex, we utilize state-of-the-art diffusion analysis techniques, including a high angular resolution diffusion imaging (HARDI) acquisition with a higher order modeling approach to enhance detail to auditory microstructure (Zanin et al., 2019). In particular, we employed multi-compartment modeling techniques including DTI with free-water elimination (fwe-DTI) and NODDI, which are made possible with a multi-shell (or multi-b-value) acquisition, and this provides a way to characterize tissue microstructure properties and reduce partial volume effects that arise from the mixing of distinct tissue compartments. We also use multi-tensor modeling of white matter fibers to improve the accuracy of fiber tracking in areas with crossing fibers, which are especially challenging in the brainstem. While the complexity and fine structure of some parts of the auditory pathway still remain beyond what may be resolved by MRI, these diffusion-derived metrics may nevertheless provide a coarse-grain representation of the major features of an individual’s auditory pathway, which are useful for establishing population-level growth models across neurodevelopment. To illustrate this point, a recent study by (Vos et al., 2015) showed DTI was sufficient to reconstruct the auditory nerve projections in a sample of adults with unilateral hearing loss; however, multi-shell dMRI tractography provided enhanced sensitivity to morphological alterations in the affected auditory nerve over single-shell methods (Ryan P Cabeen et al., 2018a). Such models can help expand our understanding of the basic neuroscience of the auditory system, and also provide a resource for potential clinical applications in diagnosis, treatment planning, and evaluation of hearing disorders.

The primary goal of this study was to elucidate the magnitude and timing of auditory pathway maturation using advanced computational neuroimaging methods applied in a large neuroimaging dataset of typically developing children (N=105) from infancy through 18 years old. We developed a novel tractography analysis pipeline to characterize the white matter bundles that constitute the ascending auditory pathway from the vestibulocochlear nerve in the brainstem to the auditory cortex. Growth models were applied to quantitative dMRI metrics derived from DTI and NODDI to characterize the magnitude and timing of microstructural features that contribute to white matter maturation of the bilateral brainstem and subcortical auditory pathways. In order to test the hypothesis that auditory pathway maturation proceeds along the inferior-to-superior axis, we use an along-tract analytical approach to characterize spatial microstructural patterns of white matter maturation along the length of each fiber bundle.

## 2. Methods

### 2.1 Subjects

Cross-sectional neuroimaging data from 105 unrelated typically developing infants, children and adolescents between 0.1 and 18.8 years of age (*M* ± *SD* = 7.8 ± 4.9 years, 56 female) were included in the present study (**Figure 1**). Children were scanned at Cincinnati Children’s Hospital Medical Center (CCHMC) as a part of the Cincinnati MR Imaging of NeuroDevelopment (C-MIND) study (Holland et al., 2015), which is publicly available at https://nda.nih.gov/edit_collection.html?id=2329. The goals of C-MIND are to characterize normative brain-behavior development and develop standardized pediatric neuroimaging protocols for typically developing children. Typically developing children were recruited through the CCHMC and were representative of the racial, ethnic and gender composition of the US population. Participants were included in the study if they or a first degree relative had a negative history of neurological or psychiatric disease, were born term (gestational age between 37 and 42 weeks), had a body mass index between the 10^th^ and 90^th^ percentile for their age and sex, and had a normal neurological exam (Kaiser et al., 2015). Subjects with abnormal MRI findings or MRI contraindications, such as orthodontic braces or metallic implants, were excluded from the study. Parent/guardian consent was obtained for all subjects, subject consent was obtained for individuals over 17 years of age, and subject assent was obtained for individuals less than 17 years of age and procedures were approved by the Institutional Review Board of CCHMC.

### 2.2 Image acquisition

Neuroimaging data was acquired on a 3T Philips Achieva scanner with a 32-channel head coil. For each subject, multi-shell diffusion weighted images (DWI) were collected across 2 scans with the following acquisition parameters: voxel size=2 mm isotropic, acquisition matrix=112×109, field of view (FOV)=224×224×120, flip angle (FA)=90° and 61 noncollinear gradient-encoding directions with 7 non-diffusion weighted (b0) images interspersed throughout each scan and averaged into a single volume. Differences in the acquisition between the 2 scans include: *Scan 1*: b=1000 s/mm^2^; repetition time (TR)/echo time (TE)=6614/81 ms; *Scan 2*: b=3000 s/mm^2^; TR/TE=8112/104 ms. T1-weighted (T1w) images were used as an anatomical reference and were acquired using a turbo-field echo magnetization-prepared rapid acquisition with gradient echo (MP-RAGE) protocol with the following parameters: voxel size=1 mm isotropic, FOV=256×224×160, FA=8°, TR/TE=8.1/3.7 ms, and inversion time (TI)=939 ms.

### 2.3 Data preprocessing

dMRI data was analyzed using Quantitative Imaging Toolkit (QIT) (R. P. Cabeen et al., 2018). Each scan was normalized by its respective b=0 volume to correct for differences in TR/TE.

FSL TOPUP and EDDY were used to correct for motion, eddy-current, and susceptibility induced geometric distortion (Andersson and Sotiropoulos, 2016) and the brain was extracted using FSL BET (Smith, 2002).. The diffusion data was spatially normalized using ANTs by first performing intra-subject registration between each subject’s DWI and T1-weighted MRIs (Avants et al., 2008). DTI-TK (Zhang et al., 2007) was then used to generate population-averaged templates of parameter maps, and templates are subsequently aligned to atlas space (Zhang et al., 2011) using the Illinois Institute of Technology (IIT) Human Brain Atlas template (Zhang and Arfanakis, 2018).

### 2.4 Diffusion modeling

Diffusion modeling was performed using the Quantitative Imaging Toolkit (QIT) (Ryan P Cabeen et al., 2018b). The parameters fractional anisotropy (FA), axial diffusivity (AD), radial diffusivity (RD) and mean diffusivity (MD) were obtained from free-water eliminated diffusion tensor imaging (fwe-DTI) (Pasternak et al., 2009) using iterative least squares optimization (Hoy et al., 2014) from the DWI acquired with b=1000 s/mm^2^. fwe-DTI has been shown to improve tissue specificity by reducing partial volume effects introduced by extracellular free water (Metzler-Baddeley et al., 2012). The neurite orientation dispersion and density imaging (NODDI) parameters neurite density index (NDI) and orientation dispersion index (ODI) (Zhang et al., 2012) were calculated using the spherical mean technique (Cabeen et al., 2019) from the concatenated DWI scans (b=1000, 3000 s/mm^2^). NODDI fixes the intrinsic diffusivity to a biologically plausible value *d_||_*= 1.7 μm^2^/ms. However, recent studies have shown shown that the default value is suboptimal in infant brains (Guerrero et al., 2019). Within white matter, (Guerrero et al., 2019) showed the optimal *d_||_*is 1.5 μm^2^/ms within a cohorts of infants less than 1 year of age. In the present dataset, the default *d_||_* was optimal in white matter of children older than 1 year of age (Supplemental Figure 1). Because a small fraction of children between 6 months and 1 year of age were included in the present study (n=8), the default *d_||_* was considered acceptable for the calculation of NODDI measures. Intra-voxel multi-tensor models to be used for tractography were calculated using FSL BEDPOSTX (Behrens et al., 2007; Jenkinson et al., 2012).

### 2.5 Tractography of the auditory pathway

We used an atlas-based streamline tractography approach to model the auditory pathway (**Figure 2**). Our modeling approach separated the auditory pathway into two distinct segments, (1) a lower brainstem pathway that was constrained to connect fibers from the internal auditory canal (IAC) to the inferior colliculus (IC), and (2) an upper pathway constrained to connect the IC to auditory cortex (AC) in Heschl’s gyrus of the superior gyrus of the temporal lobe (**Figure 1B**). We chose the split at the IC as it provided an unambiguous point of termination at each sub-pathway, and the while there are indeed intermediate synaptic connections along the auditory pathway (at the cochlear nucleus, the superior olivary complex, and the medial geniculate nucleus), the region boundaries of those intermediate nuclei do not have sufficient T1w or T2w contrast to be clearly distinguished. However, in reconstructing connections between the IAC and IC, we found the pathway trajectory traverses these intermediate structures, as indicated by 7T fMRI atlas data provided by (García-Gomar et al., 2019); hence, we adopted an approach where these intermediate points are accordingly included in the bundle model in a data-driven manner through tractography (**Figure 2C, D**). We manually labeled IAC, IC, and AC regions-of-interest (ROIs) in the IIT template (Zhang and Arfanakis, 2018). The boundaries of the AC drawn on the IIT template were defined using the anatomical landmarks described by (Desikan et al., 2006). The deformable registration algorithm in DTI-TK was used to align these ROIs to native space in each individual, and we applied a hybrid tractography algorithm (Cabeen and Toga, 2020) with the multi-tensor models to reconstruct each bundle with the following parameters: a minimum volume fraction of 0.01, maximum turning angle of 65 degrees, five seeds per voxel, a step size of 1.0 mm, and trilinear interpolation of fiber orientations. The hybrid tractography algorithm is an approach inspired by reinforcement learning to better resolve crossing fiber regions. First, probabilistic tractography is performed using the above tracking parameters, then a single dominant fiber orientation is inferred in each voxel from the most likely probabilistic bundle fiber orientation, with spatial regularization using a Markov Random Field. Subsequently deterministic streamline tractography is applied to the inferred dominant fiber orientation field. This process is described in more detail by (Cabeen and Toga, 2020), including an evaluation study of the acoustic radiation that shows this provides increased reliability and topographic regularity.

**Figure 2:**
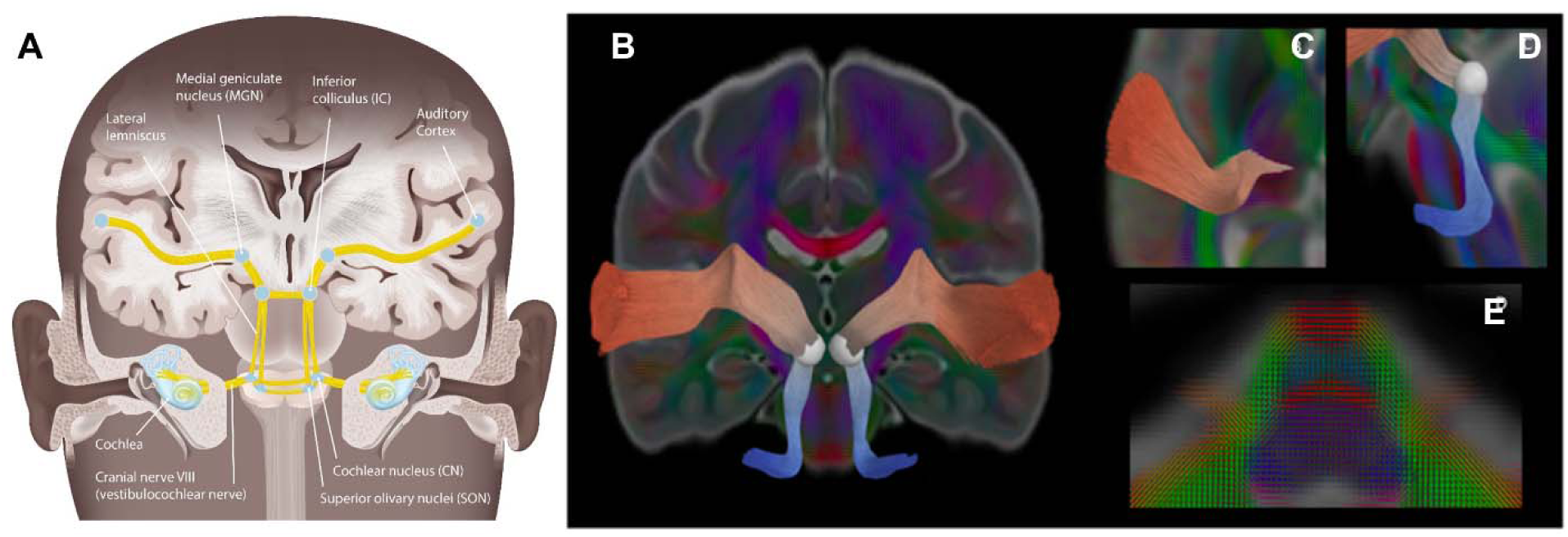
Auditory pathway anatomy. (A) Schematic of the ascending auditory pathway. (B) Tractography of the lower (blue) and upper (red) auditory pathways are shown on a coronal slice of a subject FA map overlaid with intra-voxel multi-tensor models. (C) An axial slice shows the upper auditory pathway projects through the medial geniculate nucleus of the thalamus. (D) A sagittal slice shows the auditory pathway from the cochlear nerve to the inferior colliculus (white sphere). (E) An axial slice shows the complex fiber orientations at the level of the pons.

### 2.6 Whole- and along-tract processing

The average microstructural properties within the whole lower and upper auditory pathways of the left and right hemispheres were calculated using a weighted averaging approach to avoid partial volume effects due to neighboring tissue types within the brainstem. Each estimated streamline had vertices spaces 4 mm apart and the diffusion metrics were averaged across all vertices for all streamlines. Therefore, regions of the tract with the most streamlines are weighed more heavily than regions with fewer streamlines.

We further characterized the bundles using an along tract approach, which can reveal more anatomically-specific features than available from whole bundle analysis. The along-tract technique was based on methods used in (Colby et al., 2012). First, a single prototypical “centroid” curve was estimated for each bundle from a population-averaged template using the sparse closest point transform (SCPT) that identifies the curve with the minimum mean-symmetrized Hausdorff distance to the others (Cabeen et al., 2021). The mean symmetrized Hausdorff distance has been shown to outperform other distance measures when estimating the closest point transform in tractography (Cabeen et al., 2021). The centroid curve was then resampled with distinct equidistant segments spaced every 4 mm. The template prototype curve was deformed to subject space using the transform previously computed with DTI-TK and the cross-sectional bundle vertices were labelled according to its optimal corresponding template segment label. The average NODDI and DTI measures were then computed for each group of vertices that match the centroid vertices. To account for potential misalignment in the bundle matching step, we applied along-tract smoothing using a Laplacian filter with λ = 0.3 and five iterations.

### 2.7 Statistical analysis

Statistical analyses were carried out with R version 3.1.2. The relationship between auditory tract diffusion parameters and age were assessed using the stats package (v3.6.2) with the following models: (a) a univariate linear model of the form: metric = *B_0_* + *B_1_*\*age, where *B_0_* is the y-intercept and *B_1_* is the aging coefficient or (b) a Brody growth curve implemented with nonlinear least squares regression using for the form: metric = α - (α *-* β)e^-*k*\*age^, where α is the asymptote, β is the y-intercept, and *k* is the growth rate. The best fit model for each diffusion parameter with each auditory tract and along-tract vertex was assessed using Bayesian information criteria (BIC). For models best fit with a Brody growth curve, the age at 90% of the estimated asymptotic value (age_90α_) was computed and bootstrap resampling was performed with 10,000 iterations to obtain the standard error around the estimated coefficients (boot v1.3-28). The estimated age at 90% of the exponential model plateau has been used previously to reflect the timing of developmental plateau in previous studies of white matter maturation (Chen et al., 2016; Lebel et al., 2008; Lynch et al., 2020). Lateralization in the developmental timing of auditory pathway maturation with diffusion parameters was assessed by computing the developmental laterality index (dLI) at each point along the auditory pathway: dLI = (*X_I_* - *X_C_*)/(*X_I_* + *X_C_*) where *X_I_* and *X_C_*reflect age_90α_ at corresponding points on the ipsilateral and contralateral tract, respectively. The dLI represents the relative difference in age estimates and ranges from −1 to 1, where positive values denote regions where age_90α_ is older in the ipsilateral compared to the contralateral side, while negative values denote regions where age_90α_ is younger in the ipsilateral compared to the contralateral side.

## 3. Results

### 3.1. Mean fwe-DTI and NODDI metric differences with age

The fwe-DTI metrics AD, RD and MD showed non-linear age-related differences in the bilateral upper and lower auditory pathways that were best expressed using a Brody growth curve (**Table 1**). The left upper auditory pathway also showed an exponential relationship with age using the fwe-DTI parameter FA and NODDI parameter NDI. Age-related variance in FA and NDI were best explained with a linear model for the left lower, right lower and right upper auditory pathways (**Table 2**). In these pathways, FA and NDI were significantly and positively associated with age. Age was not significantly associated with ODI in the lower (left: *p*=.09; right: *p*=.17) or upper (left: *p*=.91; right: *p*=.45) auditory pathways. **Figure 3** shows the age-related changes to fwe-DTI and NODDI parameter values averaged across the whole bilateral upper and lower auditory pathways. Per exponential model fittings, AD plateaus earliest within the right lower (age_90α_=3.15 years, 95% CI [2.04, 6.82]), left upper (age_90α_=5.72 years [4.63, 7.49]) and right upper (age_90α_=6.54 years [5.01, 9.40]) auditory pathways. RD reaches plateaus latest within the right lower (age_90α_=5.14 years [3.27, 11.93]) and upper (age_90α_=10.89 years [8.18, 16.29]) tracts, while NDI plateaus latest within the left upper tract (age_90α_=18.28 years [12.85, 33.36]). The estimated age_90α_ within the left lower tract is approximately equal across parameters with exponential fits (RD: 4.40 years [3.19, 7.03], MD: 4.58 years [3.32, 7.40], AD: 4.83 years [3.34, 8.70]). Overall, the left and right lower tracts plateau earlier across parameter values (M±SD=4.34±.70 years) compared to the left and right upper tracts (9.35±4.21 years). For MD and RD, the dLI did not significantly differ from zero, indicative of similar age_90α_ between the left and right lower tracts (RD: dLI=-.08, MD: dLI=.07), while a leftward developmental lateralization was observed in upper tracts, where age_90α_ was later in left hemisphere fibers compared to the right (RD: dLI=-.25, MD: dLI=-.20). Conversely, rightward developmental lateralization of AD was observed where age_90α_ was older in the left compared to right lower tract (dLI=.21), while no appreciable interhemispheric differences in AD age_90α_ were observed in the upper auditory pathway (dLI=-.07) (**Table 1**).

**Figure 3:**
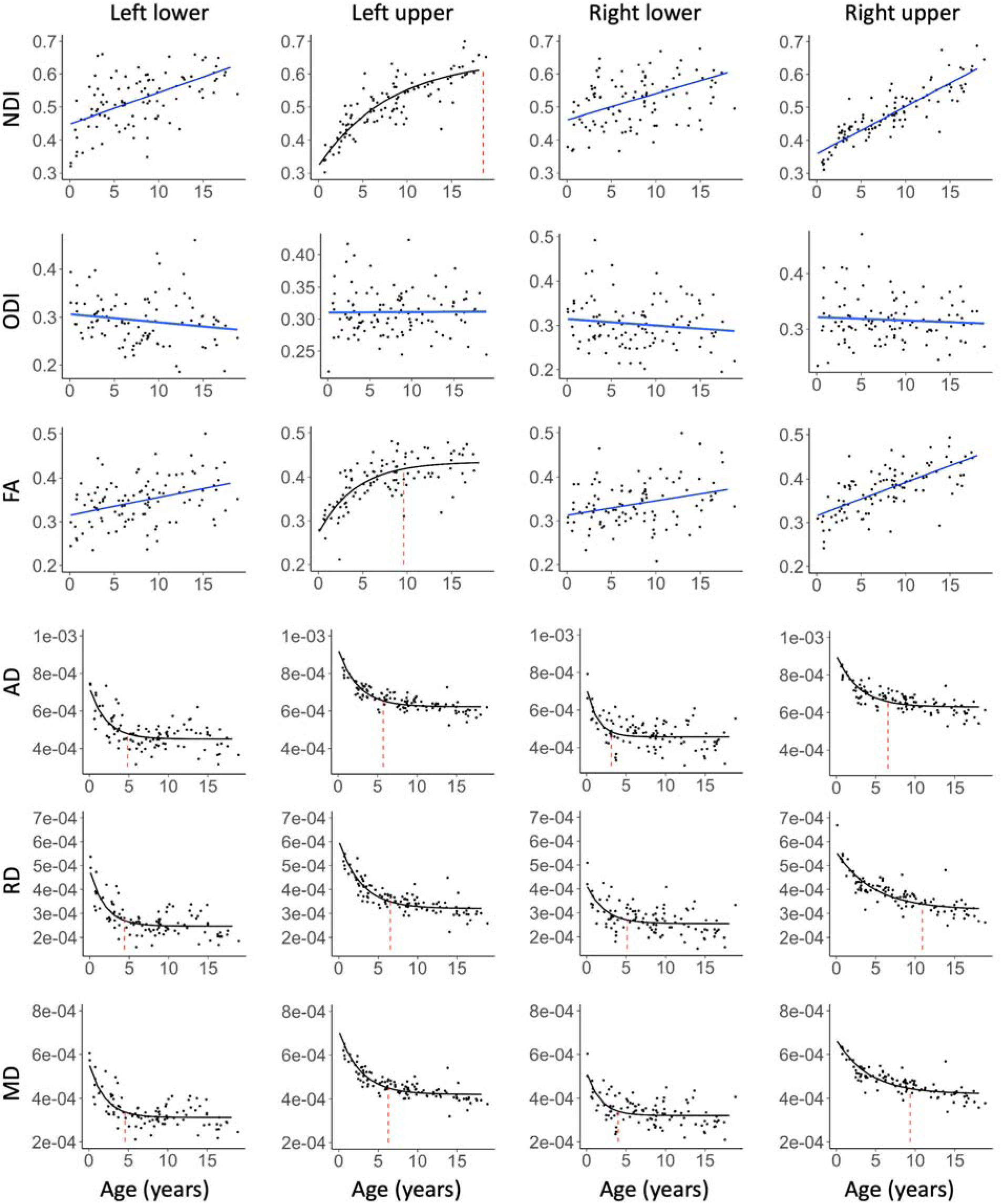
Age-related changes in auditory pathway microstructure are shown for the left and right upper and lower auditory pathways for DTI parameters (FA, AD, RD and MD) and NODDI (NDI and ODI). The fitted line for each parameter and tract reflects the best fit model, where blue denotes linear regression and black denotes exponential changes. For models that demonstrate exponential age-related changes, the age at 90% of the asymptotic value is shown with a dashed red line.

**Table 1:** Growth model parameters for auditory pathway microstructure. Fitted Brody growth curve model coefficients (coef) and standard error (SE) for the asymptote (α), growth rate (*k*) and y-intercept (*t_0_*) are shown for each metric across each auditory pathway. The y-intercept represents the estimated parameter at age=0 years. Bilateral lower tract and right upper tract FA and NDI were best fit with a linear model and growth coefficients are not provided. The age at 90% of α were calculated for tracts fit with a Brody growth curve.

| Hemisphere | Tract | Metric | $\alpha$ | | $k$ | | $t_0$ | | Age (years) |
| --- | --- | --- | --- | --- | --- | --- | --- | --- | --- |
|  |  |  | coef | SE | coef | SE | coef | SE |  |
| Left | Lower | FA | -- | -- | -- | -- | -- | -- | -- |
| | | AD | $4.50 \times 10^{-4}$ | $9.67 \times 10^{-6}$ | 0.477 | 0.112 | $7.22 \times 10^{-4}$ | $3.72 \times 10^{-5}$ | 4.83 |
| | | RD | $2.46 \times 10^{-4}$ | $6.71 \times 10^{-6}$ | 0.524 | 0.104 | $4.80 \times 10^{-4}$ | $2.75 \times 10^{-5}$ | 4.40 |
| | | MD | $3.14 \times 10^{-4}$ | $7.26 \times 10^{-6}$ | 0.503 | 0.101 | $5.60 \times 10^{-4}$ | $2.93 \times 10^{-5}$ | 4.58 |
|  |  | NDI | -- | -- | -- | -- | -- | -- | -- |
|  | Upper | FA | 0.444 | 0.014 | 0.205 | 0.053 | 0.277 | 0.018 | 11.22 |
| | | AD | $6.22 \times 10^{-4}$ | $6.01 \times 10^{-6}$ | 0.402 | 0.048 | $9.29 \times 10^{-4}$ | $2.37 \times 10^{-5}$ | 5.72 |
| | | RD | $3.20 \times 10^{-4}$ | $6.20 \times 10^{-6}$ | 0.352 | 0.042 | $6.04 \times 10^{-4}$ | $2.06 \times 10^{-5}$ | 6.54 |
| | | MD | $4.21 \times 10^{-4}$ | $5.62 \times 10^{-6}$ | 0.367 | 0.040 | $7.11 \times 10^{-4}$ | $1.96 \times 10^{-5}$ | 6.27 |
|  |  | NDI | 0.646 | 0.034 | 0.126 | 0.028 | 0.321 | 0.017 | 18.28 |
| Right | Lower | FA | -- | -- | -- | -- | -- | -- | -- |
| | | AD | $4.56 \times 10^{-4}$ | $8.41 \times 10^{-6}$ | 0.732 | 0.236 | $7.16 \times 10^{-4}$ | $4.64 \times 10^{-5}$ | 3.15 |
| | | RD | $2.54 \times 10^{-4}$ | $8.25 \times 10^{-6}$ | 0.448 | 0.148 | $4.15 \times 10^{-4}$ | $2.88 \times 10^{-5}$ | 5.14 |
| | | MD | $3.23 \times 10^{-4}$ | $7.35 \times 10^{-6}$ | 0.577 | 0.175 | $5.18 \times 10^{-4}$ | $3.38 \times 10^{-5}$ | 3.99 |
|  |  | NDI | -- | -- | -- | -- | -- | -- | -- |
|  | Upper | FA | -- | -- | -- | -- | -- | -- | -- |
| | | AD | $6.29 \times 10^{-4}$ | $7.72 \times 10^{-6}$ | 0.352 | 0.054 | $9.03 \times 10^{-4}$ | $2.51 \times 10^{-5}$ | 6.54 |
| | | RD | $3.14 \times 10^{-4}$ | $1.12 \times 10^{-5}$ | 0.211 | 0.035 | $5.55 \times 10^{-4}$ | $1.73 \times 10^{-5}$ | 10.89 |
| | | MD | $4.20 \times 10^{-4}$ | $9.06 \times 10^{-6}$ | 0.246 | 0.037 | $6.66 \times 10^{-4}$ | $1.82 \times 10^{-5}$ | 9.37 |
|  |  | NDI | -- | -- | -- | -- | -- | -- | -- |

**Table 2:** Linear model parameters for auditory pathway metrics that undergo linear age-related alterations. The fitted slope (*B*), standard error (*SE*) and t-value (*t*) for the main effect of age are shown for each metric across each auditory pathway. \**p*<.05; \*\**p*<.01; \*\*\**p*<.001

| Hemisphere | Tract | Metric | *B* | *SE* | *t* |
| --- | --- | --- | --- | --- | --- |
| Left | Lower | FA | 0.0041 | 0.0009 | 4.10*** |
|  |  | NDI | 0.0095 | 0.0015 | 6.43*** |
| Right | Lower | FA | 0.0031 | 0.0011 | 3.05** |
|  |  | NDI | 0.0079 | 0.0015 | 5.11*** |
|  | Upper | FA | 0.0078 | 0.0008 | 10.20*** |
|  |  | NDI | 0.014 | 0.0008 | 18.42*** |

### 3.2. Developmental rate and timing of tract-specific diffusion changes

Along-tract analyses demonstrate the maturational timing for each metric was not uniform (**Figure 3**). In accordance with our whole tract findings, the Brody growth curve provided the best fit model for all points along the bilateral auditory pathway for AD, RD and MD and the left upper auditory pathway for NDI. While the Brody growth curve provided the best fit model to describe age-related changes to whole left upper auditory tract FA, age-related FA changes along the most superior fibers in the superficial white matter of the left upper auditory tract were best explained with a linear model (**Figure 5**). The regions that reached age_90α_ latest across diffusion parameters include the brainstem corresponding to the inferior colliculus and superficial white matter underlying the auditory cortex (**Figure 4**). For diffusion parameters within the auditory pathway that showed nonlinear age-related changes, the standard error of the growth rate was highest in the most inferior fibers that correspond to the vestibulocochlear projections into the brainstem (**Figure 6**). Similarly, vestibulocochlear projections into the brainstem also showed the largest standard error for the estimated slope for parameters that were best fit with a linear model (**Figure 7**). **Figure 8** demonstrates the regional developmental lateralization of diffusion parameter maturation within the auditory pathway. Along-tract analyses reveal increased rightward lateralization of age_90α_ for RD and MD at the level of the inferior colliculus and the superficial white matter near the superior temporal gyrus.

**Figure 4:**
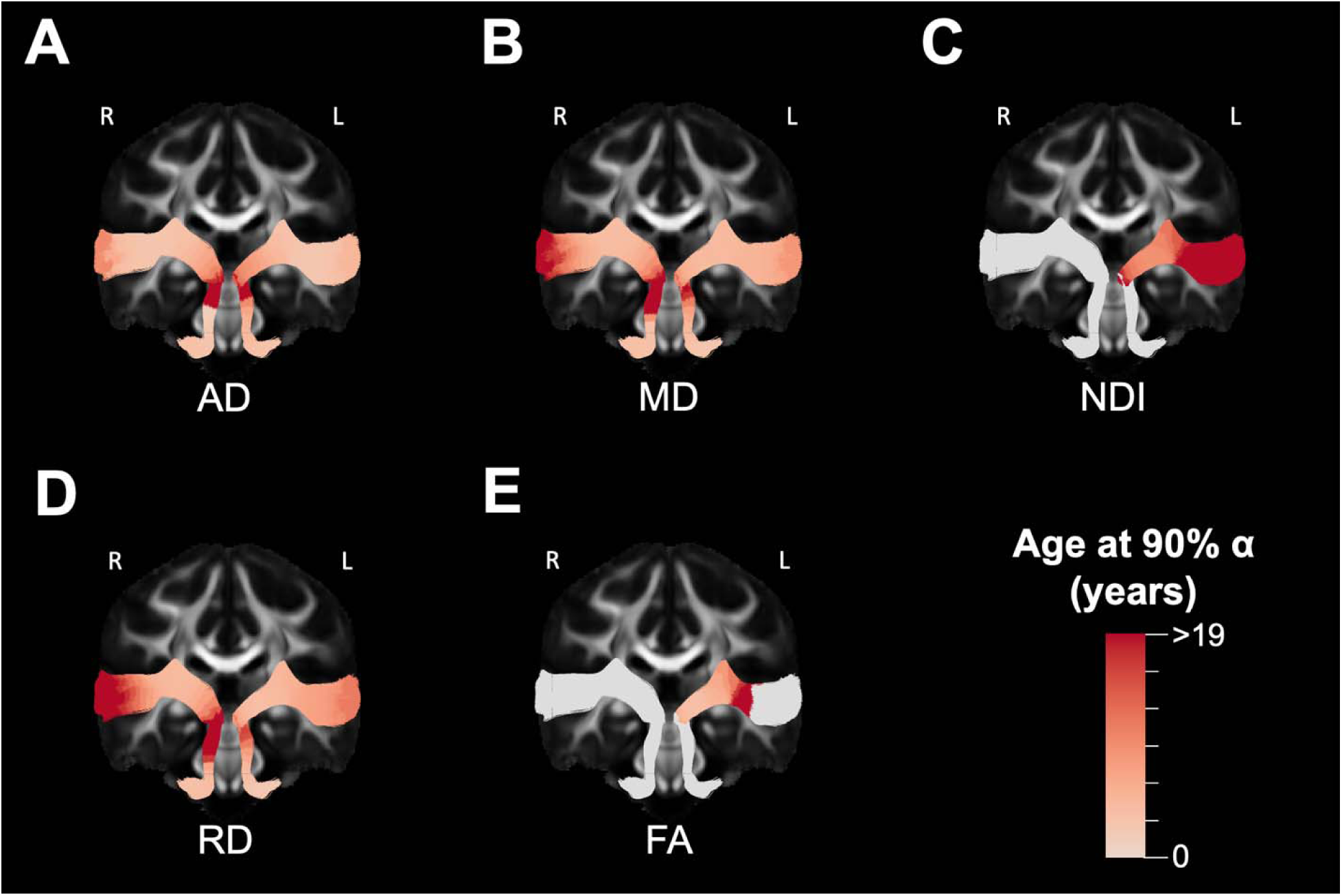
Regional variation in the developmental timing of diffusion microstructure within the auditory pathway. The age at 90% of the asymptotic value of the fitted Brody growth curve for each point along the auditory pathway are shown for (A) AD, (B) MD, (C) NDI, (D) RD and (E) FA. Regions shaded in gray denote points where a growth curve did not provide the best fit model using BIC. L – left, R – right.

**Figure 5:**
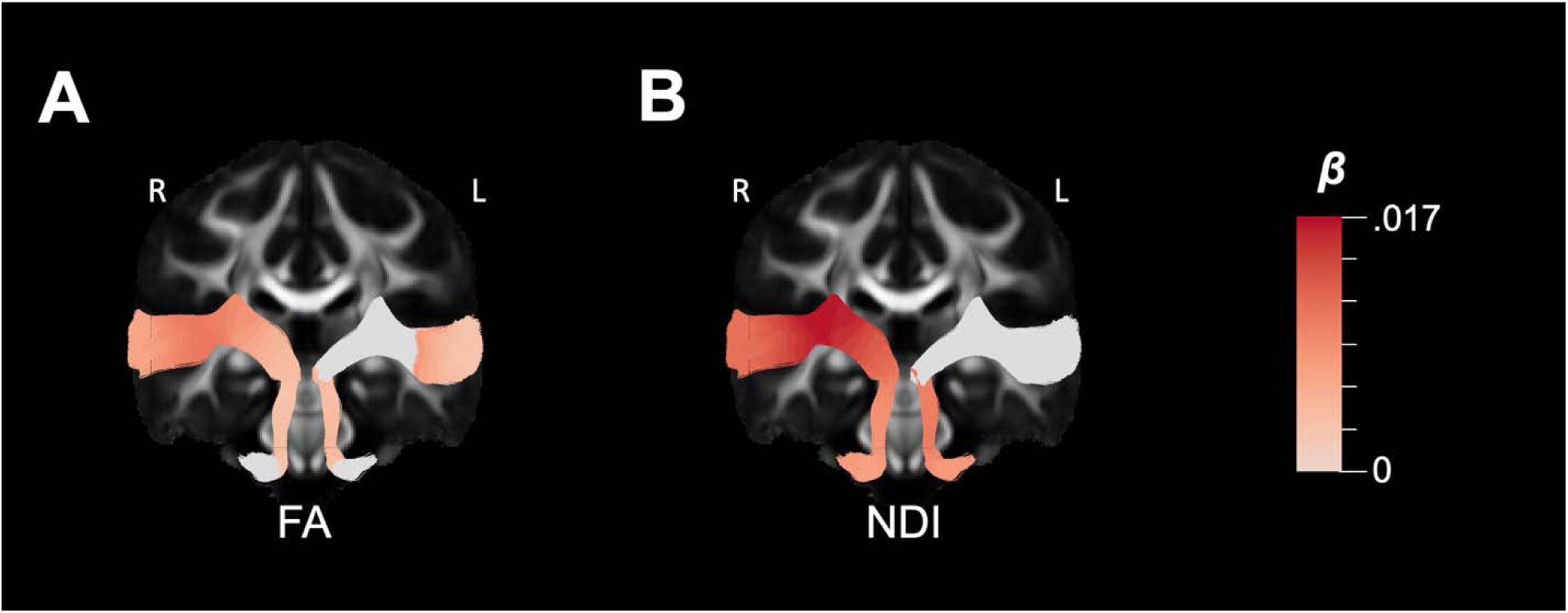
Regional linear maturation of microstructural parameters within the auditory pathway. The estimate slope (β) is provided at points along the auditory pathway for (A) FA and (B) NDI where the linear model provide the best fit model. Regions shaded in gray denote points where the linear model did not provide the best fit usin BIC. Note that FA in the bilateral vestibulocochlear nerve was not significantly associated with age using either a linear model or brody growth curve. L – left, R – right.

**Figure 6:**
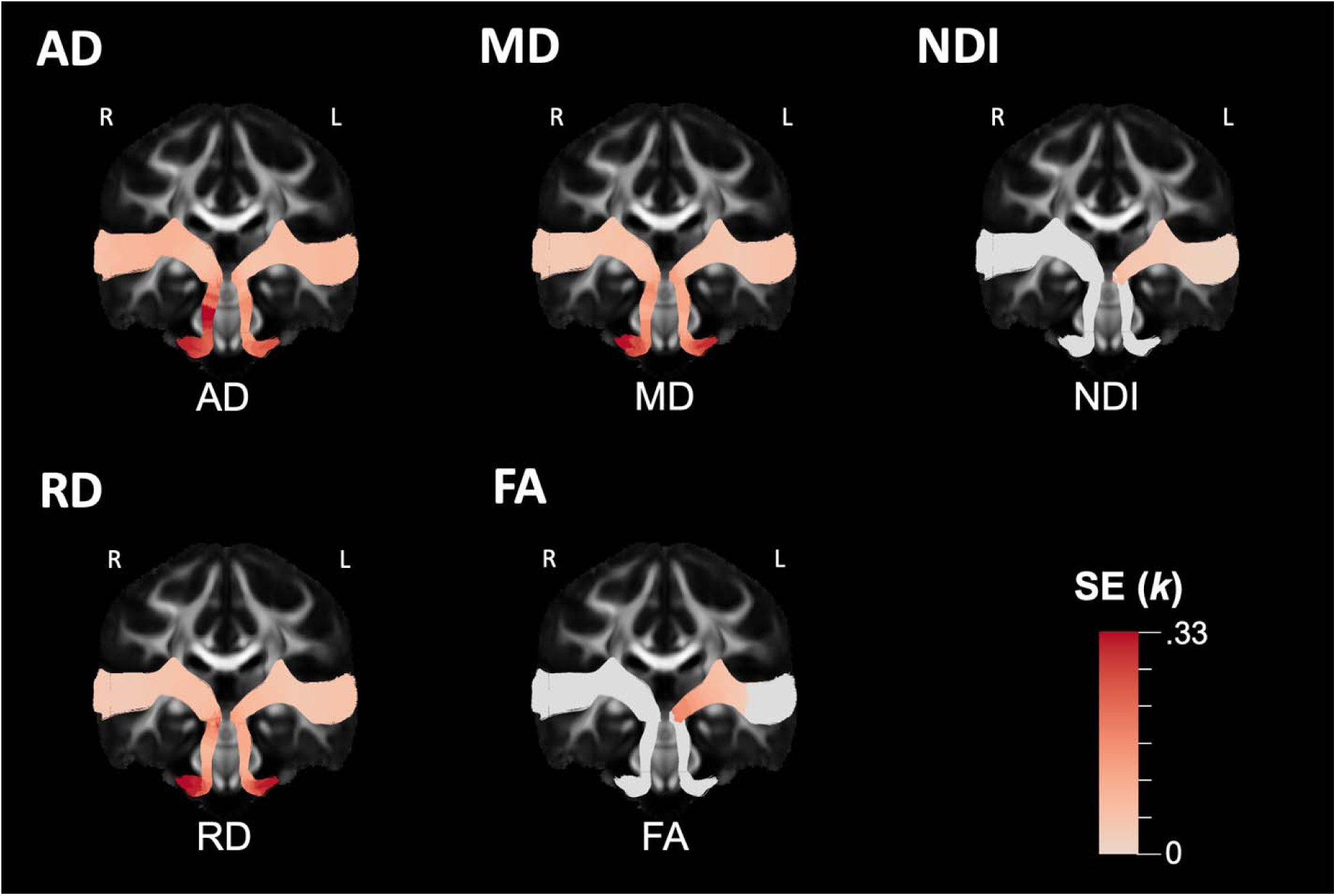
Exponential growth curve error estimates for regional auditory pathway microstructural maturation. Th standard error (SE) of the exponential growth rate (*k*) is provided for parameters along the auditory pathwa obtained through bootstrapping with replacement. Regions shaded in gray denote points where the growth curve di not provide the best fit model. The SE is provided for the same regions shown in Figure 4. L – left, R – right.

**Figure 7:**
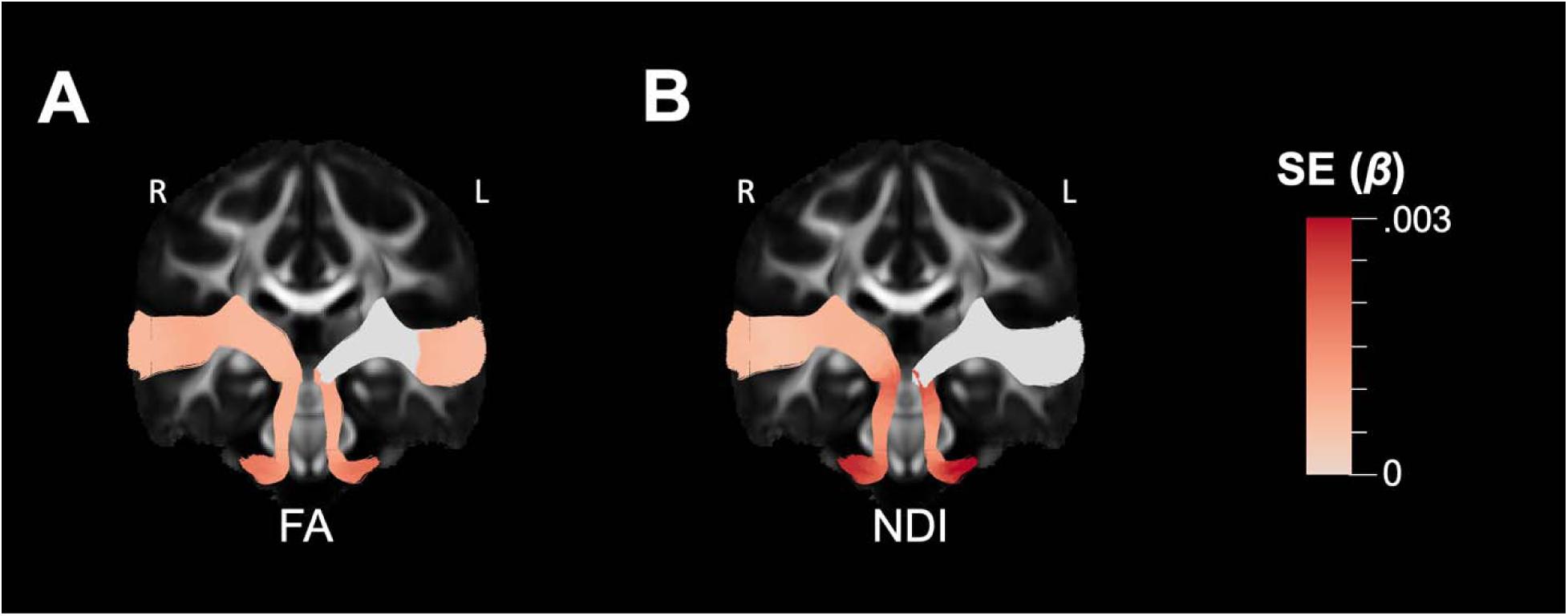
Standard error (SE) for the slope (β) estimated with a linear model for diffusion parameters at points along the auditory pathway. Regions in gray denote points where the linear model did not provide the best fit model. The SE is provided for the same regions shown in Figure 5. L – left, R – right.

**Figure 8:**
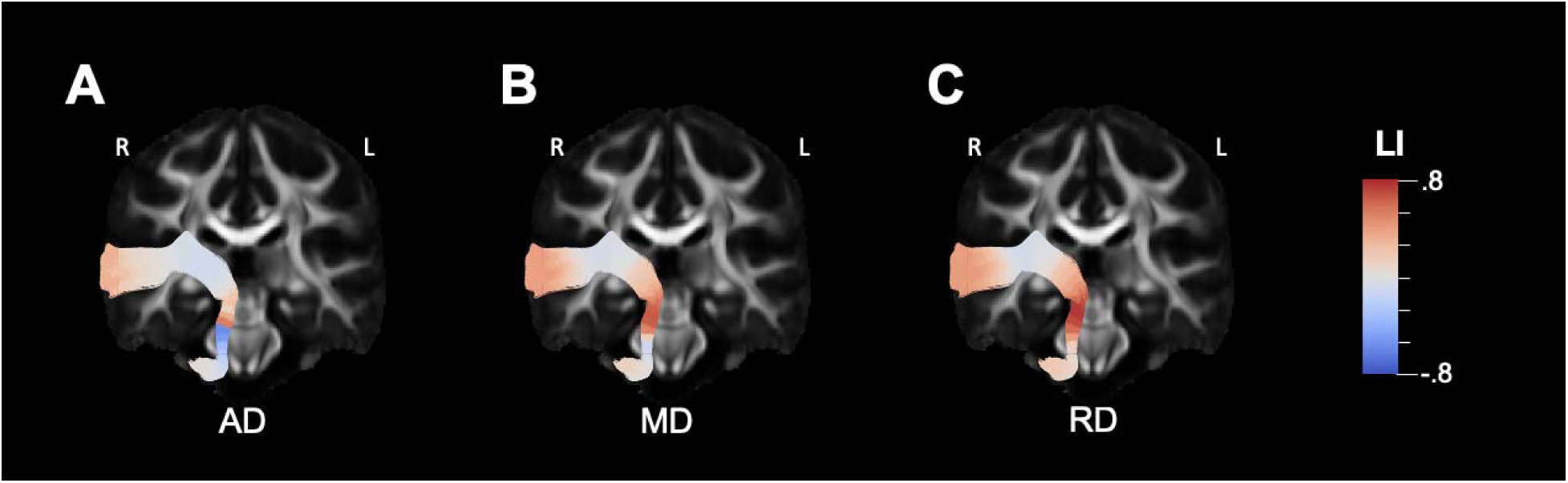
Regional lateralization of microstructural maturation within the auditory pathway. The developmental laterality index (dLI) for the age at 90% of α estimated using a Brody growth curve is provided for (A) AD, (B) MD and (C) RD. Red (positive dLI) denotes regions that develop *later* on the right side, while blue (negative dLI) denotes regions that mature *earlier* on the right side. Regions denoted as gray do not have a strongly preferre lateralization. L – left, R – right.

## 4. Discussion

This paper is the first to characterize the development of the auditory pathway of typically developing children from infancy through adolescence using diffusion neuroimaging. We have identified spatially varying patterns of white matter microstructural development from the brainstem to the subcortical white matter of the auditory cortex that differed across NODDI and fwe-DTI metrics. Our whole- and along-tract findings support previous histological studies that show a general inferior-to-superior pattern of auditory pathway maturation. Furthermore, we found the left auditory pathway reaches the estimated developmental asymptote earlier than the right hemisphere, which may be an important precursor to the development of left-lateralized language acquisition in development. Together, our findings contribute to a growing body of work that shows white matter development of auditory structures is a dynamic and heterogeneous process.

In the present study, we found AD, RD and MD are associated with rapid and nonlinear age-related decreases from infancy through adolescence, while FA and NDI increase with age and follow a more protracted developmental timeline. While evidence of auditory pathway diffusion changes across development are limited, our findings are in line with the vast majority of cross-sectional and longitudinal studies of global white matter development that show FA increases, while RD and MD decrease with age (Krogsrud et al., 2016; Lebel and Beaulieu, 2011; Lynch et al., 2020; Moura et al., 2016; Pohl et al., 2016; Scherf et al., 2014). Non-linear decreases in RD and MD likely reflect increased myelination during auditory pathway maturation. Our findings also agree with recent studies that demonstrate positive associations between NDI and age in white matter across childhood (Genc et al., 2017; Lynch et al., 2020; Mah et al., 2017). It is possible that non-linear age-related trajectories in NDI and FA were not observed due to the protracted developmental time course of these metrics, where the age range sampled was too young to detect the inflection point and asymptote of the growth curve. Therefore, the linear and protracted increase in FA and NDI likely reflect axonal maturational processes that extends well past adolescence (Bartzokis et al., 2010). This is corroborated by previous lifespan and longitudinal dMRI studies that show white matter development continues into the third decade of life (Lebel et al., 2012; Lebel and Beaulieu, 2011; Yeatman et al., 2012).

There was converging evidence across MD, AD and RD that show an inferior-to-superior spatial age-related pattern within the auditory pathway, where brainstem regions developed earlier than the thalamocortical projections to the primary auditory cortex. Within the lower auditory pathway from the vestibulocochlear nerve to the IC, AD, RD and MD reached adult levels by 5 years of age, with the earliest age estimates observed in the cranial nerves. Furthermore, the lack of a significant relationship between FA and age in the most distal portion of the lower auditory tract corresponding to the vestibulocochlear nerve suggests that developmental alterations to cochlear nerve FA was complete by the early postnatal period. Histological studies have similarly shown that rapid embryonic myelination of the cochlear nerve occurs prenatally during the second and third trimesters (Moore et al., 1997; Moore and Linthicum, 2001), while maturation of the brainstem pathways continue through the early postnatal period (Moore and Linthicum, 2007). The lower brainstem, which receives primary sensory information from the cochlear nerve, is important for the establishment of auditory perception and is highly conserved across vertebrate species (Lipovsek and Wingate, 2018). Dendritic arborization and myelination of brainstem axons within the lateral lemniscus and trapezoid body commence during the third trimester and reach adult-like levels by 6 to 12 months of age (Moore et al., 1995), which coincides with the first physiological and behavioral responses to external acoustic stimuli (Birnholz and Benacerraf, 1983; Kuhlman et al., 1988). Fibers that project from the IC to the MGN undergo the most rapid maturation during the postnatal period (Yakovlev and Lecours, 1967), which corresponds to the time when phonemic discrimination is observed in infants (Trehub, 1973). In opposition to the available evidence, we found across metrics using dMRI that the auditory pathway that projects through the IC continued to develop well past adolescence. It is unclear what the source of the discrepancy is between our findings and those found with histology, but it is possible that our findings may have been contaminated by the microstructure of neighboring structures given the small size of the IC.

Contrary to the early maturation of the brainstem auditory pathways, we found the upper auditory pathway reached asymptotic values between 5 and 11 years of age for AD, RD and MD and between 11 and 18 years for FA and NDI in the left hemisphere. For all diffusion parameters, the late maturation was driven predominantly by portions of the superficial white matter of the acoustic radiation adjacent to the auditory cortex that, for some metrics, has yet to reach adult levels by 19 years of age. Immature thalamocortical projection axons that have restricted terminations in the auditory cortical layers become apparent by 12 months of age (Moore and Guan, 2001) and proceed to myelinate into early childhood. In line with our findings, the most protracted developmental time course for the auditory pathway occurs in the intracortical projections to adjacent auditory regions within superficial white matter, where they reach adult-like levels by late childhood (Moore and Guan, 2001). Due to the increased computational capacity of intra- and inter-cortical connectivity in the auditory cortex, experience-dependent refinement of advanced auditory processing abilities also extend into adolescence (Litovsky, 2015). For example, age-dependent improvements to sound localization (Kaga, 1992), speech-in-noise discrimination (Elliott, 1979) and the identification degraded sounds (Eisenberg et al., 2000; H. et al., 2000) are observed in late childhood. Notably, the protracted structural and functional maturation of the auditory system in concert with multiple sensory and cognitive systems appear to play a crucial role in the development of speech articulation and perception, which continue to improve well into adolescence (Ross et al., 2011; Sutherland et al., 2012).

While the present study represents an important step towards characterizing normative white matter maturation within the auditory pathway, several limitations should be considered. The dMRI acquisition was collected with multiple gradient strengths to enable multi-compartment modeling; however, each shell was acquired with a different TR and TE, which may introduce a modeling bias. To overcome this issue, we normalized each diffusion scan with the non-diffusion-weighted volume of the corresponding scan to mitigate the issues introduced with differing acquisition parameters, as used in previous research (Genc et al., 2017; Lynch et al., 2020). However, results from this study should be replicated in another pediatric dataset with more similar TR/TE settings among diffusion acquisition shells. It is also possible that the spatial resolution used in the present study (2 mm isotropic) is not sufficiently high enough to capture the intricacies of the brainstem white matter pathways. Previous studies have demonstrated comparable brainstem tractography performance at spatial resolutions of 1 mm and 2 mm isotropic; however, brainstem tractography performed at the less clinically feasible .33 mm isotropic resolution provides enhanced visibility into the fine structural components of the brainstem (Ford et al., 2013). Additionally, several diffusion metrics did not reach asymptotic values and appeared to continue maturing past the age range sampled. Nonlinear fits highly depend on the sampled age range and different conclusions regarding the developmental timing of white matter maturation depend on the fitting approach and age range sampled (Lebel et al., 2019b; LeWinn et al., 2017). Therefore, a broader age range should be considered to improve our estimates of the maturational timing of auditory pathway microstructure, especially because it is known that myelination is decades-long process (Bartzokis et al., 2010). The data available for this study was cross-sectional in nature and therefore age-related trends do not reflect within-subject changes over time and inferences about white matter maturation should be interpreted with caution. Furthermore, we were unable to compare our results of auditory pathway microstructural development with functional improvements to hearing ability. Therefore, future studies should consider a longitudinal study design with functional auditory tasks to better understand the relationship between the structural and functional components of auditory development. Although, the children are considered typically developing without any clinically noted language acquisition difficulties or documented hearing loss, there were no formal audiograms obtained in the study subjects. Future studies should prospectively obtain audiograms and tests of auditory processing to confirm typical developmental processes. Additionally, inclusion of structural MRI contrasts capable of measuring specific white matter processes (i.e., myelination) should be considered, such as magnetization transfer.

To our knowledge, the present study is the first to characterize the maturational trajectory of the ascending auditory pathway in vivo from the brainstem to the auditory cortex from infancy through adolescence in a large, typically developing cohort. Utilizing along-tract analyses to gain granular insight into spatial maturational patterns of diffusion microstructure, we show brainstem regions develop than subcortical regions, and left hemisphere structures develop earlier than right. The results from study can be used as a comparative template to distinguish normative and atypical auditory pathway development. Furthermore, the proposed tractography technique will enable other researchers to delineate the fine auditory structures of the brainstem can be used broadly to study children born with abnormal auditory patterns, such as congenital or acquired sensorineural hearing loss.

## CRediT Author Statement

**Kirsten M. Lynch:** Methodology, Formal analysis, Investigation, Writing – original draft, Writing – review and editing, Visualization. **Stefanie C. Bodison**: Conceptualization, Methodology, Writing – original draft preparation, Writing – review & editing. **Ryan P. Cabeen**: Conceptualization, Methodology, Software, Writing – review and editing, Visualization. **Arthur W. Toga:** Conceptualization, Resources, Supervision, Funding acquisition. **Courtney C. J. Voelker**: Conceptualization, Methodology, Writing – original draft preparation, Writing – review & editing, Visualization, Funding acquisition.

## Data availability

All data used in this study is available through the Cincinnati MR Imaging of Neurodevelopment (C-MIND) study in the NIMH Data Archive (NDA) (https://nda.nih.gov/edit_collection.html?id=2329). Data processing modules used in the present study with the Quantitative Imaging Toolkit (QIT) can be downloaded at https://cabeen.io/qitwiki/. The tractography workflow used to extract the ascending auditory pathway, including region of interest masks and bundle definitions, can be found here: https://github.com/cabeen/auditory-pathway

## Acknowledgements

We acknowledge the contributions of Dr. Laurel Fisher, who brought valuable insight, enthusiasm, and kindness to the project and team, and Caroline O’Driscoll for her assistance with illustrations used in the present study.

