## Supplementary Figure 1 for "The spatial organization of ascending auditory pathway microstructural maturation from infancy through adolescence using a novel fiber tracking approach"

**Supplementary Material**

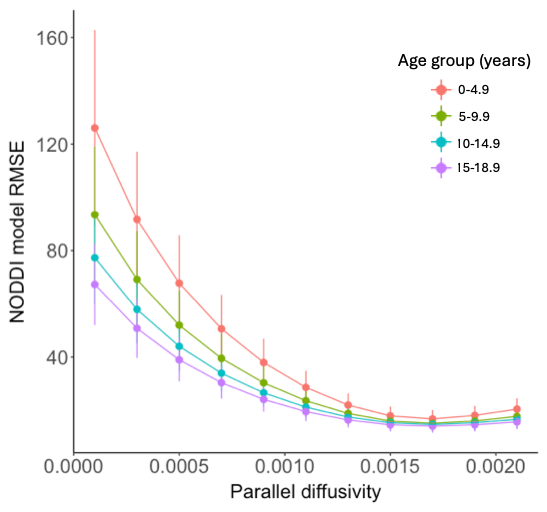

**Supplemental Figure 1**. The influence of intrinsic parallel diffusivity fixation (d_||_, μm^2^/ms) on NODDI model root mean square error (RMSE) in the white matter in 5-year binned groups. Across age groups, the optimal d_||_ = .0017 μm^2^/ms.
